# Uncovering DUB Selectivity Through Ion-Mobility-Based Assessment of Ubiquitin Chain Isomers

**DOI:** 10.1101/2023.10.11.561976

**Authors:** Elizaveta I. Shestoperova, Eric R. Strieter

## Abstract

Ubiquitination is a reversible posttranslational modification that maintains cellular homeostasis and regulates protein turnover. Deubiquitinases (DUBs) are a large family of proteases that catalyze the removal of ubiquitin (Ub) along with the dismantling and editing of Ub chains. Assessing the activity and selectivity of DUBs is critical for defining physiological function. Despite numerous methods for evaluating DUB activity, none are capable of assessing activity and selectivity in the context of multicomponent mixtures of native, unlabeled ubiquitin conjugates. Here we report on an ion mobility (IM)-based approach for measuring DUB selectivity in the context of unlabeled mixtures of Ub chains. We show that IM-MS can be used to assess the selectivity of DUBs in a time-dependent manner. Moreover, using the branched Ub chain selective DUB UCH37/UCHL5 along with a mixture of Ub trimers, a strong preference for branched Ub trimers bearing K6 and K48 linkages is revealed. Our results demonstrate that IM coupled with mass spectrometry (IM-MS) is a powerful method for evaluating DUB selectivity under conditions more physiologically relevant than single component mixtures.

## INTRODUCTION

Ubiquitination is a posttranslational modification that strongly regulates protein turnover in eukaryotic cells.^1,2^ It is well established that ubiquitination triggers various outcomes in cells, including protein proteasomal degradation, regulation of gene expression, and apoptosis, showing that ubiquitination can store and transmit information.^3–5^ The ubiquitination mechanism is now well understood and consists of a three-step enzymatic cascade, including E1 ubiquitin (Ub)-activating, E2 Ub-conjugating, and E3 Ub-ligating enzymes.^6–8^Besides monoubiquitination of the substrate at single or multiple sites, Ub can be attached as a polymeric chain that is formed through a covalent bond between the ε-amino group of the lysine (K6, K11, K27, K29, K33, K48, K63) or N-terminal methionine (M1) of one Ub monomer and carboxyl terminus of another.^9^ These polyUb chains vary in length, linkage, and degree of branching (i.e., how many Ub subunits are modified at more than one site with other Ub molecules), forming a complex Ub code that determines the cellular fate of ubiquitinated proteins.^10–12^

The Ub code is tightly regulated by a family of proteolytic enzymes referred to as deubiquitinases (DUBs)^13^. Among the ∼100 human DUBs, only a fraction has been characterized with regards to substrate selectivity.^14^ In most cases, selectivity—which is often considered a proxy for biological function—is assessed using homogenous mixtures of synthetic or semi-synthetic Ub conjugates^15–21^. These efforts have led to several important discoveries, namely that some DUBs selectively target certain ubiquitinated proteins and chain types.^14,22,23^ Polyubiquitinated proteins, however, coexist in cells and there are a dearth of studies examining the impact of natural competition on selectivity of DUBs.^24^ A recent report used a set of neutron-encoded di-Ub derivatives to evaluate the selectivity of DUBs against all eight linkage types simultaneously. Distinct masses for each di-Ub were obtained using synthetic Ub monomers bearing distinct patterns of heavy isotopes.^25^ Although this study represents a significant advance, it will be challenging to extend this method to longer chains and use cell lysates. Additional methods are therefore needed to assess DUB selectivity in the context of heterogenous Ub chain mixtures.

Here we report on the use of ion mobility mass spectrometry (IM-MS) to inform on the selectivity of DUBs using complex mixtures of Ub chains. We show that tri-Ub isomers with different linkages and chain topologies exhibit unique collision induced unfolding (CIU) fingerprints. Instead of selecting a specific collision voltage (CV), we found that the entire CIU fingerprint can be used to deconvolute CIU patterns within a complex mixture of isomers. We exploit this attribute to assess the selectivity of DUBs using a mixture of tri-Ubs. Our results demonstrate that when coupled with a deconvolution algorithm, IM-MS can be a powerful method for evaluating the selectivity of DUBs in complex mixtures.

## EXPERIMENTAL METHODS

### Protein expression and purification

Wild-type Ub along with lysine-to-arginine variants were expressed in E. coli Rosetta 2(DE3)pLysS cells and purified by perchloric acid precipitation, following procedure adapted from ref.^26^All enzymes were expressed in Rosetta 2(DE3)pLysS or BL21(DE3)pLysS *E.coli* cells, and further purified with affinity chromatography as previously described.^27–33^ The detailed description on the protein expression and purification protocols is provided in the Supporting Information.

### Generation of native Ub chains

*K48-linked tri-Ub:* 2 mM Ub, 1 uM E1, and 10 uM UBE2R1 were mixed in reaction buffer A (20 mM ATP, 10 mM MgCl^2^, 40 mM Tris-HCl pH 7.5, 50 mM NaCl, and 1.5 mM DTT) and incubated for 6 h at 37°C. *K63-linked tri-Ub:* 1 mM Ub, 1.5 uM E1, and 0.75 uM Ubc13-Mms2 were mixed in reaction buffer A and incubated at 37°C for 9 h. *K6-linked tri-Ub:* 2 mM Ub, 0.5 uM E1, 10 uM UbcH7, and 1uM NleL were mixed in reaction buffer A. 5 uM OTUB1 and 3 uM AMSH were added after 3 h of incubation and the mixture was left overnight at 37°C.

*K6/K48 branched triUb:* 2 mM Ub K6R/K48R, 1 mM Ub D77, 0.5 uM E1, 10 uM UbcH7, 1 uM NleL, 3 uM AMSH were mixed in reaction buffer A and incubated overnight at 37

°C. *K48/K63-branched tri-Ub:* 2 mM Ub K48R/K63R, 1 mM Ub D77, 0.5 uM E1, 10 uM UBE2R1, 1uM UBE2N/UBE2V2 were mixed in reaction buffer A and incubated overnight at 37 °C. *K11/K48-branched tri-Ub:* 2 mM Ub K11R/K48R 1 mM Ub D77, 0.5 uM E1, 10 uM UBE2R1, 20 uM UBE2S-UBD, 1uM UBE2N/UBE2V2 were mixed in reaction buffer A and incubated overnight at 37 °C. For all branched trimers, the enzymatic reaction mixture was further treated with 0.5 uM Yuh1 at room temperature overnight to remove the C-terminal D77 residue from the proximal Ub. All trimers (homotypic and heterotypic branched) were purified using size exclusion chromatography (Superdex 75) with 150 mM ammonium acetate pH 6.8. Concentrations of all trimers were measured by BCA assay.

### Debranching assays

*Gel-based debranching assay of single isomers:* 0.25 mg/mL of single tri-Ub isomers were treated with 1 uM AMSH, 1 uM OTUB1, 3 uM UCH37, or 5 uM UCH37·Rpn13 in assay buffer (50 mM HEPES pH 7.5, 50 mM NaCl, and 2 mM DTT). The reactions were quenched at specific time points with 6xLaemmli loading buffer before SDS-PAGE analysis. *For IM-MS analysis, the samples were prepared as described:* the mixtures containing 0.25 mg/mL of each tri-Ub isomer were treated with DUBs using the same conditions as gel-based debranching assay. The reactions were quenched with 100 mM ammonium acetate pH 4.4 at specific time points to precipitate the enzymes and exchanged into a buffer containing 150 mM ammonium acetate Ph 6.8 prior to IM-MS analysis.

### ESI-IM-MS analysis

All experiments were performed on Waters Synapt G2 HDMS mass spectrometer in native conditions using NanoLockSpray ion source for offline nanoESI. Each Ub trimer was diluted to a final concentration of 10 uM in 150 mM ammonium acetate pH 6.8. The IM separation was performed at capillary voltage – 1.0 kV, source temperature – 20°C, sampling cone – 20 V, extraction cone – 4.4 V, trap gas flow – 2 mL/min, helium cell gas flow – 180 mL/min, IMS gas flow – 60 mL/min. TWIMS parameters were adjusted to receive the most efficient separation between Ub trimers at highest available wave height – 40 V. All CIU analyses were performed by increasing the trap collision voltage in a stepwise manner from 4−80 V in 10 V increments. All CIU fingerprints displayed in the paper are the result of three replicates. The complex mixtures containing multiple tri-Ub isomers were obtained by mixing defined amounts of single isomers at the equal molar ratios.

### Data analysis

The initial processing of IM data was performed using MassLynx MS Software (Waters Corporation, USA). IM spectra were extracted using TWIM extract.^34^ The CIU heat-maps were obtained using CIUSuite2^35^. The deconvolution of the CIU fingerprints with CIU fingerprints of single components was performed using custom python-based scripts (see Supporting Information for more details). Further data analyses were performed using the Software package OriginPro 2023 (OriginLab Corporation, USA).

## RESULTS AND DISCUSSION

### Collision-induced unfolding (CIU) of tri-Ub ions

Building on previous studies showing di-Ub isomers have unique CIU fingerprints, we wanted to determine whether unfolding patterns could be used to distinguish more complex Ub chains, including those containing different linkage compositions and chain topology.^36^ We selected a library of four tri-Ub isomers containing K6-, K48-, K63-linked linear, and K6/K48-branched trimers. These isomers were chosen based on the availability of highly specific approaches for enzymatic synthesis.^37–40^ Figure 1 demonstrates the unfolding patterns of single tri-Ubs at two major charge states (Figure S1). Collisional activation leads to sequential activation of numerous unfolded conformations in the gas phase for each isomer. However, the sheer number of gas-phase conformations makes it challenging to select specific collision voltages that are optimal for distinguishing isomers simultaneously. Thus, we reasoned it would be possible to use the whole CIU pattern for a specified isomer without selecting specific collision voltages.

**Figure 1.**
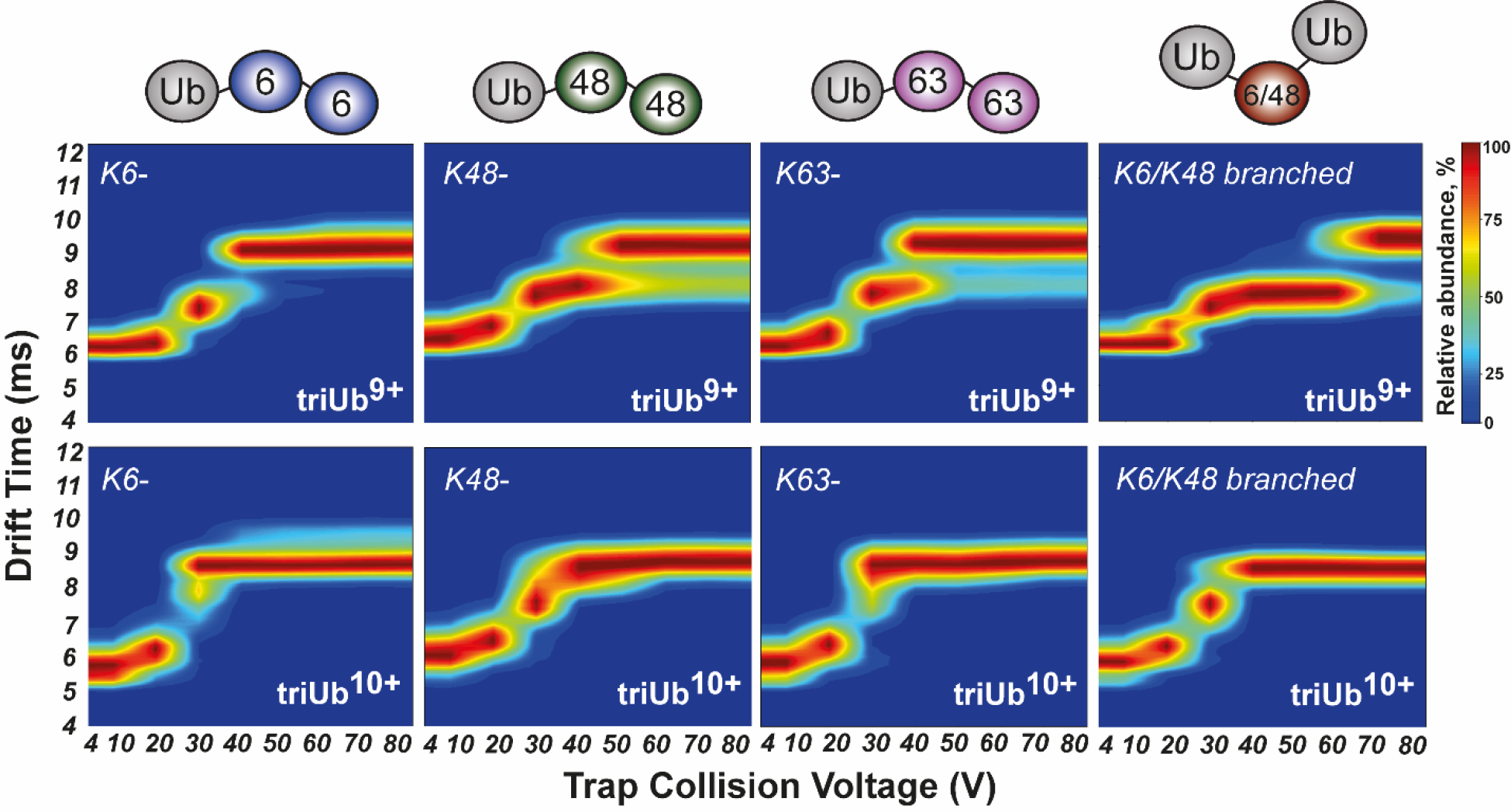
CIU fingerprints for K6-, K48-, K63-, and K6/K48 branched-linked tri-Ub (left to right) at +9 and +10 charge states (top to bottom).

### Deconvolution algorithm

We previously showed that multiple linear regression analysis could enable IM spectral deconvolution of mixtures containing di-Ub isomers. Euclidean distances were measured between spectra of a complex mixture and a linear combination of individual isomer IM spectra to assess the degree of similarity^41^. We aimed to expand the previously developed approach to perform the simultaneous fitting of all IM spectra from the CIU pattern of a complex mixture with the IM spectra from the CIU patterns of individual isomers (assuming that the contributions from each isomer to the complex mixture spectra are consistent across the whole CIU pattern). To this end, we had to modify our algorithm to include the sum of the Euclidean distances between the experimental IM spectra of a complex mixture and the linear combination of single IM spectra collected at all collision voltages. The experimental data was fitted iteratively by optimizing the weights for each isomer to minimize the deviation between the experimental and fitted spectra. A formal description of the deconvolution algorithm is shown in the Supporting Information.

We first tested our CIU fitting algorithm starting with a 1:1 mixture of unbranched K6 and branched K6/K48 tri-Ub isomers (Figure 2A). The whole CIU fingerprint deconvolution is visualized on a set of four collision voltages (20, 40, 60, and 70 V) (Figure 2B).

**Figure 2.**
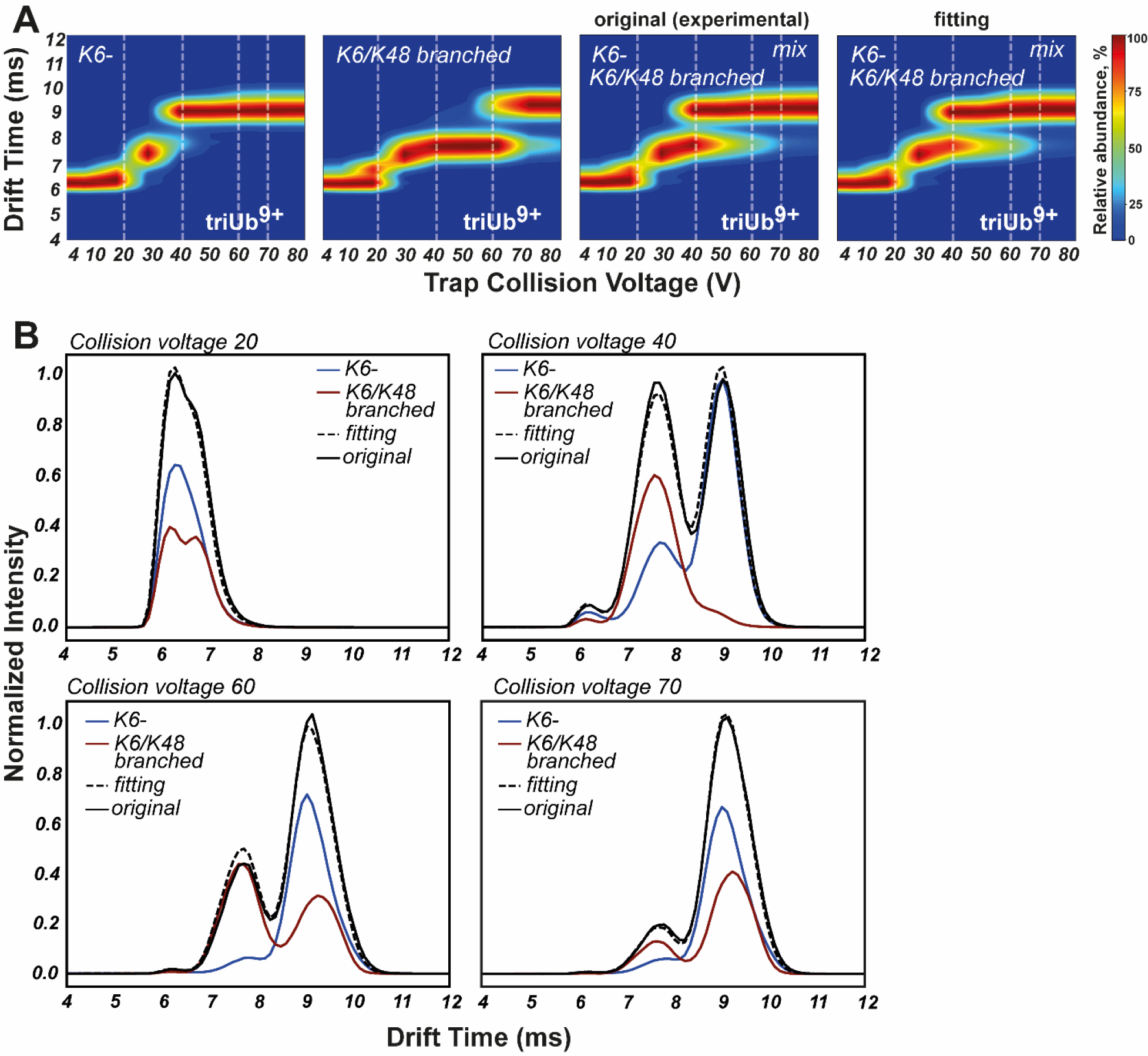
Deconvolution of a binary mixture consisting of K6 unbranched and K6/K48 branched tri-Ub. A) CIU fingerprints for K6 tri-Ub, K6/K48 branched tri-Ub, and a 1:1 mixture of K6 and K6/K48 tri-Ub. The calculated CIU fingerprint of a 1:1 binary mixture was obtained using the deconvolution algorithm described herein. CIU data was obtained under native conditions (150 mM ammonium acetate, pH 6.8) at +9 charge state. The white dash lines represent the chosen CV used for demonstration of the fitting in Figure 2B. B) Deconvolution of the IM spectra of the binary mixture with the spectra of single isomers at a specific CV. The black solid line represents the experimental IM spectra at different CVs. The black dash line represents the IM spectra of the binary mixture obtained from the fitting. The fitted CIU fingerprints represent the sum of CIU fingerprints of single isomers multiplied by corresponding relative weights calculated from the deconvolution.

Euclidean distances were measured between the IM spectra from the CIU pattern of the binary mixture (Figure 2B; black lines) and linear combinations of IM spectra of individual isomers to find the relative weights of each isomer in the mixture. The individual IM spectra at different CV were then multiplied by the calculated relative weights to form the optimal fitting of the experimental data (Figure 2B; dashed lines). As shown in Figure 2B, there is a good match between the calculated fit and the experimental data, demonstrating the deconvolution approach works for the entire CIU pattern of a binary mixture.

### Qualitative assessment of DUB selectivity

Next, we wanted to apply our deconvolution approach to more complex mixtures. Using a total of four distinct trimers, we prepared different ternary mixtures and measured the corresponding CIU patterns. Deconvolution with CIU fingerprints of all four single isomers correctly identifies the isomers present in each mixture along with the one that is absent. A cut-off value of 0.1, which was used to identify the missing isomer, is necessary to account for the experimental variability in IM data (Figure 3, Figure S2, Table S1).

**Figure 3.**
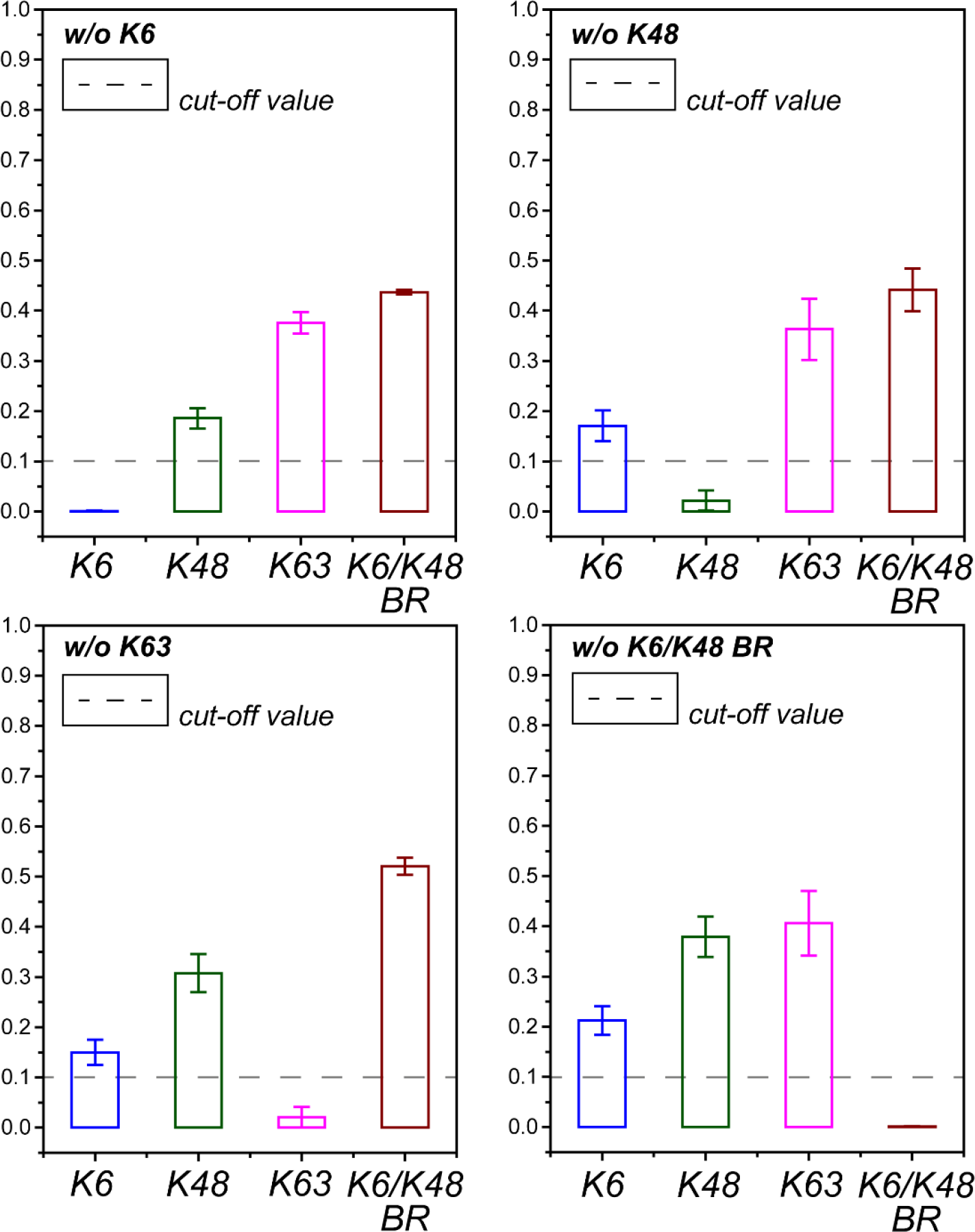
Relative weights of tri-Ub isomers in ternary mixtures obtained from deconvolution. The white dashed line represents the cut-off value below which the isomer is considered absent from the mixture. The data was averaged between three repeats within the same biological sample.

We then assessed whether deconvolution of a complex mixture can inform on the selectivity of a DUB. To this end, we used DUBs with selectivity toward specific Ub chain types, e.g., the K63-linkage specific DUB AMSH, the K48-specific DUB OTUB1, and the branched chain selective DUB UCH37/UCHL5.^40,42,43^ Reactions were performed with a quaternary mixture of Ub trimers and CIU patterns were obtained and deconvoluted to calculate the weights of each remaining isomer after complete conversion by a particular DUB. All four isomers were detected upon deconvolution of the quaternary mixture before DUB treatment. After cleavage, we obtained results consistent with the expected selectivities: K63-tri-Ub is cleaved by AMSH, K48-tri-Ub by OTUB1, and the K6/K48 branched tri-Ub is dismantled by UCH37 (Figure 4, Figure S3, Table S2). These data demonstrate that deconvolution of CIU patterns can inform on the selectivity of DUBs without using isotopically labeled Ub chains or any reporter tags.

**Figure 4.**
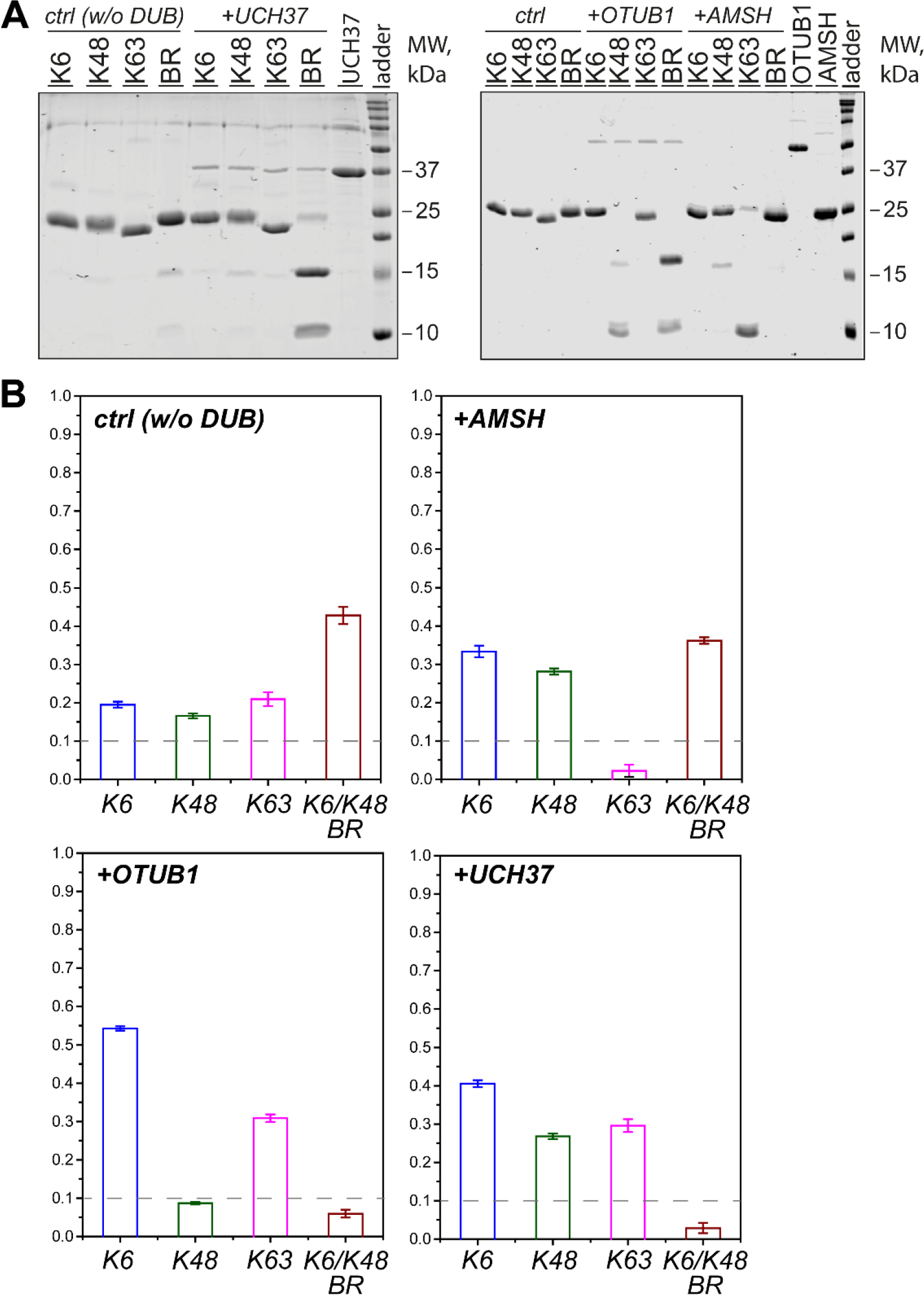
Qualitative analysis of DUB selectivity. A) Gel-based deubiquitination assays with UCH37/UCHL5 (left), OTUB1 and AMSH (right). 3 uM of UCH37/UCHL5, 1 uM OTUB1 or 1 uM AMSH was mixed 10 uM of each Ub trimer and incubated for 3 h at 37°C. B) CIU-based deubiquitination assays. Quaternary mixture containing K6-, K48-, K63-, and K6/K48-branched trimers was treated with UCH37/UCHL5, OTUB1 or AMSH. The relative weights after DUB cleavage obtained from the deconvolution algorithm are shown. The white dashed line represents the cut-off value below which the isomer is considered absent from the mixture. The data was averaged between three repeats within the same biological sample.

### Time-dependent assessment of DUB activity

Having demonstrated the use of IM-MS in examining the selectivity of DUBs on a qualitative level, we wanted to determine whether CIU-based deconvolution could be used to track time-dependent changes in Ub chain composition. To this end, we monitored the cleavage activity of UCH37 using binary, ternary, and quaternary mixtures of Ub chains. Due to the selective debranching activity of UCH37, we expected to observe selective depletion of K6/K48 tri-Ub from mixtures in which K6/K48 tri-Ub is the only branched chain present. The composition of each mixture was assessed at six (for binary and ternary) and five (for quaternary) different timepoints. As expected, we observed a gradual decrease in the molar fraction of K6/K48 tri-Ub in each of the mixtures along with a concomitant increase in the molar fractions of the unbranched chains (Figure 5, Figures S4-S5, Table S3). To further test the utility of our approach, we wanted to determine which type of branched trimer is preferred by UCH37 in the presence of competition. For these experiments, we decided to use UCH37 in complex with the proteasomal binding partner RPN13 to improve cleavage efficiency.^40^The UCH37•RPN13 complex was subjected to an equimolar mixture of three different branched trimers: K6/K48, K11/K48, and K48/K63. The reaction was monitored over time by IM-MS. The data show that the K6/K48 branched trimer is cleaved first, followed by K48/K63 and finally K11/K48 (Figure 6, Figure S6, Table S4). These results—which are consistent with previous reports with isolated branched trimers—demonstrate that UCH37 strongly prefers K6/K48 branched trimers^44^.

**Figure 5.**
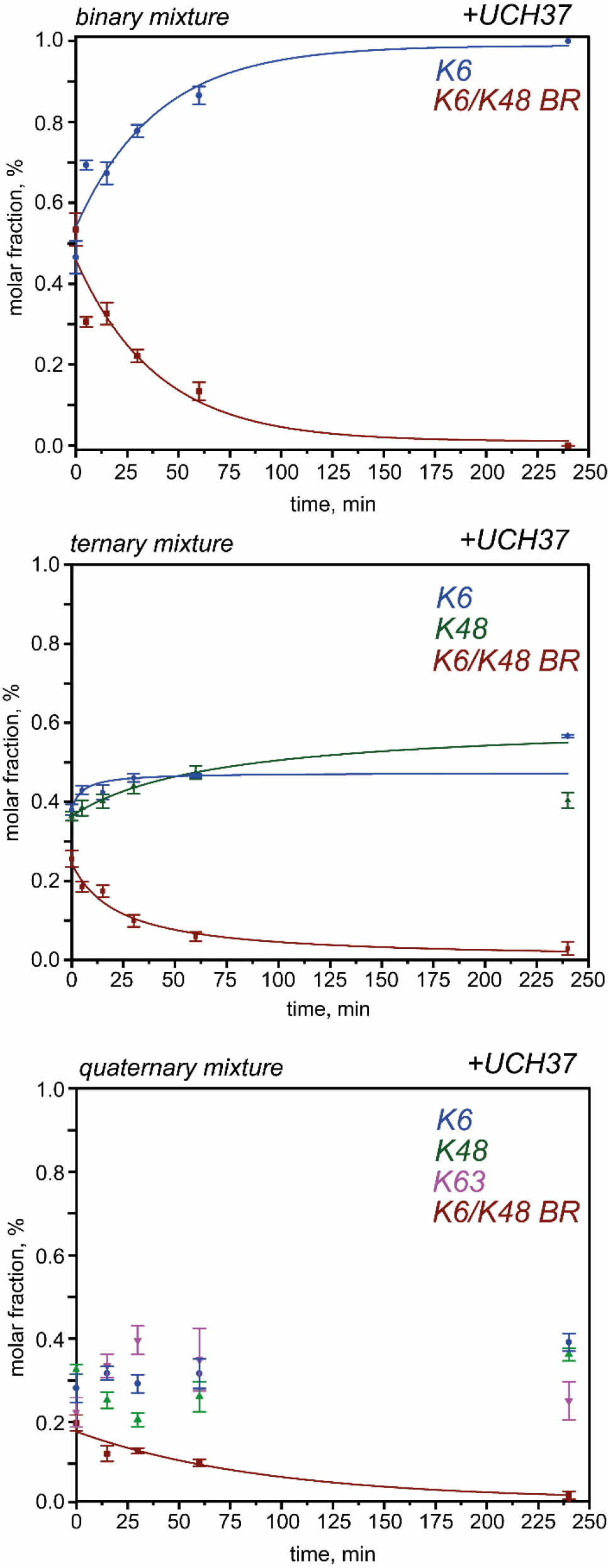
Time-course cleavage assays with UCH37/UCH5. *Top panel*: time-course analysis of a binary mixture consisting of K6-, and K6/K48-branched trimers; *middle panel:* time-course analysis of a ternary mixture consisting of K6-, K48-, and K6/K48-branched trimers; *bottom panel:* time-course analysis of a quaternary mixture consisting of K6-, K48-, K63- and K6/K48-branched trimers. The mixtures containing 10 uM of each isomer were incubated with 3 uM UCH37/UCHL5 for the indicated time points (0, 5, 15, 30, 60, 240 min for ternary and 0, 15, 30, 60, 240 min for quaternary mixtures). The CIU-based deconvolution was applied to calculate the molar fraction of single components at each time point. The calculated weights were averaged between three repeats within the same biological sample.

**Figure 6.**
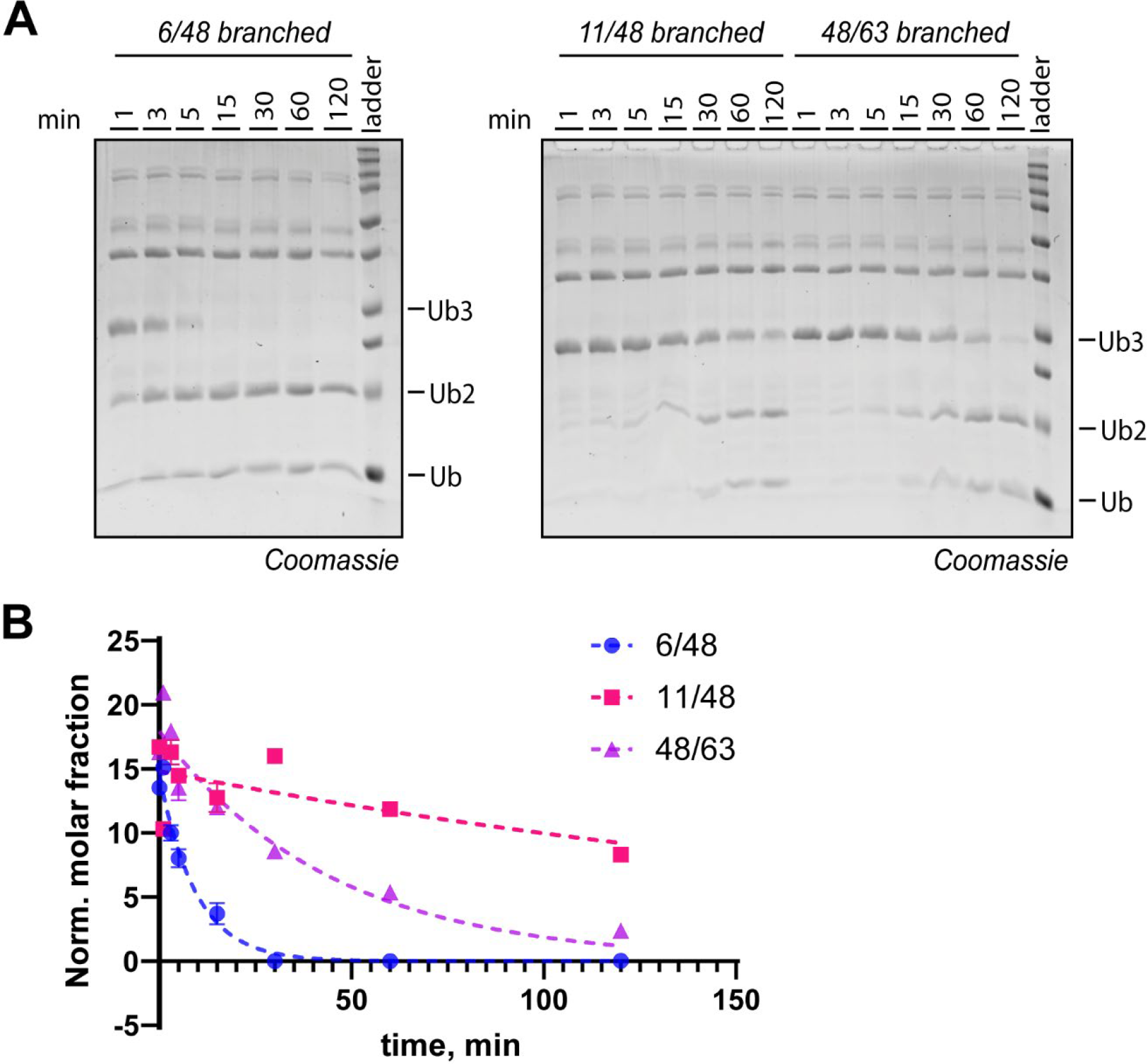
Time-course cleavage assay of branched tri-Ubs using UCH37·Rpn13. A) Gel-based cleavage assay with isolated branched tri-Ubs. 10 uM of each isomer was incubated with 5 uM of UCH37·Rpn13. The reaction was quenched at the indicated time points (0, 1, 3, 5, 15, 30, 60, 120 min). B) CIU-based deconvolution to determine molar fractions of single tri-Ubs in a complex mixture during cleavage with UCH37·Rpn13. The calculated weights were averaged between three repeats within the same biological sample.

## CONCLUSIONS

In conclusion, we developed the CIU-based deconvolution approach to assess DUB selectivity on mixtures of native isomeric Ub chains. Using the whole CIU fingerprints, multicomponent mixtures of Ub chains composed of three subunits can be deconvoluted even if there is a high degree of similarity in the unfolding patterns of the different chain types. Underscoring the utility of this approach, we show the selectivity of DUBs can be evaluated under scenarios in which multiple unlabeled Ub chains compete for recognition. This led to the finding that the proteasomal DUB UCH37 exhibits a strong preference for K6/K48 branched trimers. Thus, IM-MS has the potential to facilitate the analysis of DUBs under physiologically relevant settings.

## NOTES

The authors declare no competing financial interest(s).

## ASSOCIATED CONTENT

The Supporting Information is available. (SI S3) deconvolution algorithm for fitting of CIU data, calculation of deconvolution errors; (Table S1) calculated molar fractions of single tri-Ub isomers in different ternary (out-of-one) mixtures; (Table S2) calculated molar fractions of single tri-Ub isomers in the quaternary mixture treated with various DUBs; (Table S3) calculated molar fractions of single tri-Ub isomers in the quaternary mixture treated with UCH37 (time-dependent); (Table S4) calculated molar fractions of branched tri-Ub isomers in the ternary mixture treated with UCH37·Rpn13 (time-dependent); (Figure S1) MS1 spectra of tri-Ub isomers in native conditions; (Figure S2) demonstration of the CIU-based deconvolution stability on the example of ternary mixture lacking 6/48-branched isomer; (Figure S3) CIU fingerprints that were obtained during the qualitative assessment of DUB specificity; (Figure S4) gel-based assay of UCH37/UCHL5-dependent degradation of 6/48-branched trimer; (Figure S5) CIU fingerprints for the time-course degradation assays of binary, ternary, or quaternary mixtures with UCH37/UCHL5; (Figure S6) CIU fingerprints for the time-course degradation assays of branched trimers with UCH37·Rpn13; (SI S14-S15) extended protein expression and purification protocols.

## ACKNOWLEDGEMENTS

This work was supported by research grant R35GM149532 from the National Institutes of Health (NIH). The authors wish to thank the UMass Amherst Institute of Applied Life Sciences Mass Spectrometry Core Facility (RRID:SCR_019063) for access to the Waters Synapt G2 HDMS. We thank UMass Amherst Institute of Applied Life Sciences Mass Spectrometry Core Facility director Dr. Stephen Eyles.

## REFERENCES

(1) Komander, D.; Rape, M. The Ubiquitin Code. Annu. Rev. Biochem. 2012. 10.1146/annurev-biochem-060310-170328.

(2) Damgaard, R. B. The Ubiquitin System: From Cell Signalling to Disease Biology and New Therapeutic Opportunities. Cell Death and Differentiation. 2021. 10.1038/s41418-020-00703-w.

(3) Ciechanover, A.; Schwartz, A. L. The Ubiquitin-Proteasome Pathway: The Complexity and Myriad Functions of Proteins Death. Proceedings of the National Academy of Sciences of the United States of America. 1998. 10.1073/pnas.95.6.2727.

(4) Yao, T.; Ndoja, A. Regulation of Gene Expression by the Ubiquitin-Proteasome System. Semin. Cell Dev. Biol. 2012, 23 (5), 523–529. 10.1016/j.semcdb.2012.02.006.

(5) Vucic, D.; Dixit, V. M.; Wertz, I. E. Ubiquitylation in Apoptosis: A Post-Translational Modification at the Edge of Life and Death. Nature Reviews Molecular Cell Biology. 2011. 10.1038/nrm3143.

(6) Schulman, B. A.; Wade Harper, J. Ubiquitin-like Protein Activation by E1 Enzymes: The Apex for Downstream Signalling Pathways. Nature Reviews Molecular Cell Biology. 2009. 10.1038/nrm2673.

(7) Ye, Y.; Rape, M. Building Ubiquitin Chains: E2 Enzymes at Work. Nature Reviews Molecular Cell Biology. 2009. 10.1038/nrm2780.

(8) Buetow, L.; Huang, D. T. Structural Insights into the Catalysis and Regulation of E3 Ubiquitin Ligases. Nature Reviews Molecular Cell Biology. 2016. 10.1038/nrm.2016.91.

(9) Deol, K. K.; Lorenz, S.; Strieter, E. R. Enzymatic Logic of Ubiquitin Chain Assembly. Frontiers in Physiology. 2019. 10.3389/fphys.2019.00835.

(10) Ohtake, F.; Tsuchiya, H. The Emerging Complexity of Ubiquitin Architecture. J. Biochem. 2016. 10.1093/jb/mvw088.

(11) Akutsu, M.; Dikic, I.; Bremm, A. Ubiquitin Chain Diversity at a Glance. J. Cell Sci. 2016. 10.1242/jcs.183954.

(12) French, M. E.; Koehler, C. F.; Hunter, T. Emerging Functions of Branched Ubiquitin Chains. Cell Discov. 2021, 7 (1), 6. 10.1038/s41421-020-00237-y.

(13) Amerik, A. Y.; Hochstrasser, M. Mechanism and Function of Deubiquitinating Enzymes. Biochimica et Biophysica Acta - Molecular Cell Research. 2004. 10.1016/j.bbamcr.2004.10.003.

(14) Clague, M. J.; Urbé, S.; Komander, D. Publisher Correction: Breaking the Chains: Deubiquitylating Enzyme Specificity Begets Function. Nat. Rev. Mol. Cell Biol. 2019, 20 (5), 321–321. 10.1038/s41580-019-0112-8.

(15) El Oualid, F.; Merkx, R.; Ekkebus, R.; Hameed, D. S.; Smit, J. J.; de Jong, A.; Hilkmann, H.; Sixma, T. K.; Ovaa, H. Chemical Synthesis of Ubiquitin, Ubiquitin-Based Probes, and Diubiquitin. Angew. Chemie Int. Ed. 2010, 49 (52), 10149–10153. 10.1002/anie.201005995.

(16) Kumar, K. S. A.; Spasser, L.; Erlich, L. A.; Bavikar, S. N.; Brik, A. Total Chemical Synthesis of Di-Ubiquitin Chains. Angew. Chemie - Int. Ed. 2010. 10.1002/anie.201003763.

(17) McGouran, J. F.; Gaertner, S. R.; Altun, M.; Kramer, H. B.; Kessler, B. M. Deubiquitinating Enzyme Specificity for Ubiquitin Chain Topology Profiled by DiUbiquitin Activity Probes. Chem. Biol. 2013. 10.1016/j.chembiol.2013.10.012.

(18) Dang, L. C.; Melandri, F. D.; Stein, R. L. Kinetic and Mechanistic Studies on the Hydrolysis of Ubiquitin C-Terminal 7-Amido-4-Methylcoumarin by Deubiquitinating Enzymes. Biochemistry 1998. 10.1021/bi9723360.

(19) Ritorto, M. S.; Ewan, R.; Perez-Oliva, A. B.; Knebel, A.; Buhrlage, S. J.; Wightman, M.; Kelly, S. M.; Wood, N. T.; Virdee, S.; Gray, N. S.; Morrice, N. A.; Alessi, D. R.; Trost, M. Screening of DUB Activity and Specificity by MALDI-TOF Mass Spectrometry. Nat. Commun. 2014, 5 (1), 4763. 10.1038/ncomms5763.

(20) Mulder, M. P. C.; El Oualid, F.; Ter Beek, J.; Ovaa, H. A Native Chemical Ligation Handle That Enables the Synthesis of Advanced Activity-Based Probes: Diubiquitin as a Case Study. ChemBioChem 2014. 10.1002/cbic.201402012.

(21) Geurink, P. P.; van Tol, B. D. M.; van Dalen, D.; Brundel, P. J. G.; Mevissen, T. E. T.; Pruneda, J. N.; Elliott, P. R.; van Tilburg, G. B. A.; Komander, D.; Ovaa, H. Development of Diubiquitin-Based FRET Probes To Quantify Ubiquitin Linkage Specificity of Deubiquitinating Enzymes. ChemBioChem 2016, 17 (9), 816–820. 10.1002/cbic.201600017.

(22) Mevissen, T. E. T.; Hospenthal, M. K.; Geurink, P. P.; Elliott, P. R.; Akutsu, M.; Arnaudo, N.; Ekkebus, R.; Kulathu, Y.; Wauer, T.; El Oualid, F.; Freund, S. M. V.; Ovaa, H.; Komander, D. OTU Deubiquitinases Reveal Mechanisms of Linkage Specificity and Enable Ubiquitin Chain Restriction Analysis. Cell 2013, 154 (1), 169–184. 10.1016/j.cell.2013.05.046.

(23) Faesen, A. C.; Luna-Vargas, M. P. A.; Geurink, P. P.; Clerici, M.; Merkx, R.; Van Dijk, W. J.; Hameed, D. S.; El Oualid, F.; Ovaa, H.; Sixma, T. K. The Differential Modulation of USP Activity by Internal Regulatory Domains, Interactors and Eight Ubiquitin Chain Types. Chem. Biol. 2011. 10.1016/j.chembiol.2011.10.017.

(24) Tracz, M.; Bialek, W. Beyond K48 and K63: Non-Canonical Protein Ubiquitination. Cell. Mol. Biol. Lett. 2021, 26 (1), 1. 10.1186/s11658-020-00245-6.

(25) van Tol, B. D. M.; van Doodewaerd, B. R.; Lageveen-Kammeijer, G. S. M.; Jansen, B. C.; Talavera Ormeño, C. M. P.; Hekking, P. J. M.; Sapmaz, A.; Kim, R. Q.; Moutsiopoulou, A.; Komander, D.; Wuhrer, M.; van der Heden van Noort, G. J.; Ovaa, H.; Geurink, P. P. Neutron-Encoded Diubiquitins to Profile Linkage Selectivity of Deubiquitinating Enzymes. Nat. Commun. 2023, 14 (1), 1661. 10.1038/s41467-023-37363-6.

(26) Pickart, C. M.; Raasi, S. Controlled Synthesis of Polyubiquitin Chains. Methods in Enzymology. 2005. 10.1016/S0076-6879(05)99002-2.

(27) Bashore, C.; Dambacher, C. M.; Goodall, E. A.; Matyskiela, M. E.; Lander, G. C.; Martin, A. Ubp6 Deubiquitinase Controls Conformational Dynamics and Substrate Degradation of the 26S Proteasome. Nat. Struct. Mol. Biol. 2015. 10.1038/nsmb.3075.

(28) Michel, M. A.; Elliott, P. R.; Swatek, K. N.; Simicek, M.; Pruneda, J. N.; Wagstaff, J. L.; Freund, S. M. V.; Komander, D. Assembly and Specific Recognition of K29- and K33-Linked Polyubiquitin. Mol. Cell 2015. 10.1016/j.molcel.2015.01.042.

(29) Pham, G. H.; Rana, A. S. J. B.; Korkmaz, E. N.; Trang, V. H.; Cui, Q.; Strieter, E. R. Comparison of Native and Non-Native Ubiquitin Oligomers Reveals Analogous Structures and Reactivities. Protein Sci. 2016, 25 (2), 456–471. 10.1002/pro.2834.

(30) Trang, V. H.; Valkevich, E. M.; Minami, S.; Chen, Y. C.; Ge, Y.; Strieter, E. R. Nonenzymatic Polymerization of Ubiquitin: Single-Step Synthesis and Isolation of Discrete Ubiquitin Oligomers. Angew. Chemie - Int. Ed. 2012. 10.1002/anie.201207171.

(31) Valkevich, E. M.; Sanchez, N. A.; Ge, Y.; Strieter, E. R. Middle-Down Mass Spectrometry Enables Characterization of Branched Ubiquitin Chains. Biochemistry 2014. 10.1021/bi5006305.

(32) Bremm, A.; Freund, S. M. V.; Komander, D. Lys11-Linked Ubiquitin Chains Adopt Compact Conformations and Are Preferentially Hydrolyzed by the Deubiquitinase Cezanne. Nat. Struct. Mol. Biol. 2010. 10.1038/nsmb.1873.

(33) Du, J.; Babik, S.; Li, Y.; Deol, K. K.; Eyles, S. J.; Fejzo, J.; Tonelli, M.; Strieter, E. A Cryptic K48 Ubiquitin Chain Binding Site on UCH37 Is Required for Its Role in Proteasomal Degradation. Elife 2022, 11. 10.7554/eLife.76100.

(34) Haynes, S. E.; Polasky, D. A.; Dixit, S. M.; Majmudar, J. D.; Neeson, K.; Ruotolo, B. T.; Martin, B. R. Variable-Velocity Traveling-Wave Ion Mobility Separation Enhancing Peak Capacity for Data-Independent Acquisition Proteomics. Anal. Chem. 2017. 10.1021/acs.analchem.7b00112.

(35) Polasky, D. A.; Dixit, S. M.; Fantin, S. M.; Ruotolo, B. T. CIUSuite 2: Next-Generation Software for the Analysis of Gas-Phase Protein Unfolding Data. Anal. Chem. 2019. 10.1021/acs.analchem.8b05762.

(36) Wagner, N. D.; Clemmer, D. E.; Russell, D. H. ESI-IM-MS and Collision-Induced Unfolding That Provide Insight into the Linkage-Dependent Interfacial Interactions of Covalently Linked Diubiquitin. Anal. Chem. 2017. 10.1021/acs.analchem.7b02932.

(37) Hospenthal, M. K.; Freund, S. M. V.; Komander, D. Assembly, Analysis and Architecture of Atypical Ubiquitin Chains. Nat. Struct. Mol. Biol. 2013. 10.1038/nsmb.2547.

(38) Dong, K. C.; Helgason, E.; Yu, C.; Phu, L.; Arnott, D. P.; Bosanac, I.; Compaan, D. M.; Huang, O. W.; Fedorova, A. V.; Kirkpatrick, D. S.; Hymowitz, S. G.; Dueber, E. C. Preparation of Distinct Ubiquitin Chain Reagents of High Purity and Yield. Structure 2011. 10.1016/j.str.2011.06.010.

(39) Hofmann, R. M.; Pickart, C. M. In Vitro Assembly and Recognition of Lys-63 Polyubiquitin Chains. J. Biol. Chem. 2001. 10.1074/jbc.M103378200.

(40) Deol, K. K.; Crowe, S. O.; Du, J.; Bisbee, H. A.; Guenette, R. G.; Strieter, E. R. Proteasome-Bound UCH37/UCHL5 Debranches Ubiquitin Chains to Promote Degradation. Mol. Cell 2020, 80 (5), 796–809.e9. 10.1016/j.molcel.2020.10.017.

(41) Shestoperova, E. I.; Ivanov, D. G.; Strieter, E. R. Quantitative Analysis of Diubiquitin Isomers Using Ion Mobility Mass Spectrometry. J. Am. Soc. Mass Spectrom. 2023. 10.1021/jasms.3c00016.

(42) McCullough, J.; Clague, M. J.; Urbé, S. AMSH Is an Endosome-Associated Ubiquitin Isopeptidase. J. Cell Biol. 2004, 166 (4), 487–492. 10.1083/jcb.200401141.

(43) Edelmann, M. J.; Iphöfer, A.; Akutsu, M.; Altun, M.; di Gleria, K.; Kramer, H. B.; Fiebiger, E.; Dhe-Paganon, S.; Kessler, B. M. Structural Basis and Specificity of Human Otubain 1-Mediated Deubiquitination. Biochem. J. 2009, 418 (2), 379–390. 10.1042/BJ20081318.

(44) Song, A.; Hazlett, Z.; Abeykoon, D.; Dortch, J.; Dillon, A.; Curtiss, J.; Martinez, S. B.; Hill, C. P.; Yu, C.; Huang, L.; Fushman, D.; Cohen, R. E.; Yao, T. Branched Ubiquitin Chain Binding and Deubiquitination by Uch37 Facilitate Proteasome Clearance of Stress-Induced Inclusions. Elife 2021. 10.7554/eLife.72798.

