## Supporting Information for "Uncovering DUB Selectivity Through Ion-Mobility-Based Assessment of Ubiquitin Chain Isomers"

### TABLE OF CONTENTS


| | | |
|---|---|---|
| | Deconvolution algorithm for fitting of CIU data | S3 |
| | Calculations of errors in molar fractions of tri-Ub isomers upon CIU-based deconvolution | S3 |
| Table S1. | Molar fractions of single tri-Ub isomers in different ternary (out-of-one) mixtures | S4 |
| Table S2. | Molar fractions of single tri-Ub isomers in the quaternary mixture treated with various DUBs | S5 |
| Table S3. | Molar fractions of single tri-Ub isomers in the quaternary mixture treated with UCH37 (time-dependent) | S6 |
| Table S4. | Molar fractions of branched tri-Ub isomers in the ternary mixture treated with UCH37·Rpn13 (time-dependent) | S7 |
| Figure S1. | MS1 spectra of tri-Ub in the native state | S8 |
| Figure S2. | Demonstration of deconvolution stability | S9 |
| Figure S3. | CIU fingerprints that were obtained during the qualitative assessment of DUB specificity | S10 |
| Figure S4. | UCH37/UCHL5-dependent degradation of 6/48-branched trimer (SDS-PAGE) | S11 |
| Figure S5. | CIU fingerprints for the time-course degradation assays of binary, ternary, or quaternary mixtures with UCH37/UCHL5 | S12 |
| Figure S6. | CIU fingerprints for the time-course degradation assays of branched trimers with UCH37·Rpn13 | S13 |
| | Protein expression/purification | S14-S15 |

**Deconvolution algorithm for fitting of CIU data**

*1. Input data*

$\boldsymbol{k}$ – a set of triUb isomers

$\boldsymbol{I_k(dt, CV)}$ – CIU fingerprint for a single triUb isomer

$I_k(dt, CV)$, where dt – ATD (arrival time distribution); CV – collision voltage, CV $\in \{4, 10, \dots .80\}$

$\boldsymbol{I(dt, CV)}$ – CIU fingerprint for an unknown mixture

The goal is to find the best fitting of $I(dt, CV)$ as a linear combination of $I_k(dt, CV)$ for each CV value.

*2. Variables*

$\boldsymbol{w_k}$ – relative weight of an isomer (the same for all CV values)

$\boldsymbol{\varphi_{CV}}$ – normalized coefficient (accounts for differences of absolute intensities of ATD at different CV)

*3. Statement of problem*

For all possible collision voltages find the set of $\{w_k, \varphi_{CV}\}$ that results in the best fit of $I(dt, CV)$ CIU fingerprint as a linear combination of individual $I_K(dt, CV)$ CIU patterns, or:

$$I(dt, CV) = \varphi_{CV} \cdot \sum I_k \cdot w_k \qquad (1)$$

*4. Optimization problem*

$$G(w_k, \varphi_{CV}) := \sum_{CV} \left\| I(dt, CV) - \varphi_{CV} \sum I_k(dt, CV) \cdot w_k \right\|$$

$$G(w_k, \varphi_{cv}) \xrightarrow[\sum w_k = 1]{} min$$

**Calculations of errors in molar fractions of tri-Ub isomers upon CIU-based deconvolution**

The standard deviation for the molar fraction values was calculated using the equation:

$$SD = \sqrt{\frac{\sum_{i=1}^{n} \left(\chi_{i,} - \chi_{i,average}\right)^2}{n(n-1)}}\ , where\ n - number\ of\ the\ components\ in\ the\ mixture$$

**Table S1.** Molar fractions of single tri-Ub isomers in different ternary mixtures. Molar fraction values were obtained by deconvolution of the CIU fingerprint of different ternary mixtures with CIU fingerprints of four (K6, K48-, K63, 6/48-branched) isomers. The mean molar fraction values (<x>) were calculated from three repeats within the same biological sample. Standard deviation was calculated as shown in page S3.

| ***w/o 6/48-branched*** | | | | | ***w/o K6-*** | | | | |
|---|---|---|---|---|---|---|---|---|---|
| 6/48-branched | | | | | 6/48-branched | | | | |
| rep1 | rep2 | rep3 | <χ> | SD | rep1 | rep2 | rep3 | <χ> | SD |
| 0 | 0 | 0 | 0 | 0 | 0.444 | 0.438 | 0.429 | 0.437 | 0.004359 |
| K6 | | | | | K6 | | | | |
| rep1 | rep2 | rep3 | <χ> | SD | rep1 | rep2 | rep3 | <χ> | SD |
| 0.162 | 0.261 | 0.215 | 0.212667 | 0.028603 | 0 | 0 | 0 | 0 | 0 |
| K48 | | | | | K48 | | | | |
| rep1 | rep2 | rep3 | <χ> | SD | rep1 | rep2 | rep3 | <χ> | SD |
| 0.301 | 0.4 | 0.436 | 0.379 | 0.040361 | 0.215 | 0.147 | 0.196 | 0.186 | 0.020257 |
| K63 | | | | | K63 | | | | |
| rep1 | rep2 | rep3 | <χ> | SD | rep1 | rep2 | rep3 | <χ> | SD |
| 0.535 | 0.337 | 0.347 | 0.406333 | 0.064398 | 0.34 | 0.414 | 0.374 | 0.376 | 0.021385 |
| ***w/o K48-*** | | | | | ***w/o K63-*** | | | | |
| 6/48-branched | | | | | 6/48-branched | | | | |
| rep1 | rep2 | rep3 | <χ> | SD | rep1 | rep2 | rep3 | <χ> | SD |
| 0.391 | 0.526 | 0.409 | 0.442 | 0.04232 | 0.553 | 0.494 | 0.515 | 0.520667 | 0.017266 |
| K6 | | | | | K6 | | | | |
| rep1 | rep2 | rep3 | <χ> | SD | rep1 | rep2 | rep3 | <χ> | SD |
| 0.126 | 0.23 | 0.157 | 0.171 | 0.030827 | 0.126 | 0.123 | 0.201 | 0.15 | 0.025515 |
| K48 | | | | | K48 | | | | |
| rep1 | rep2 | rep3 | <χ> | SD | rep1 | rep2 | rep3 | <χ> | SD |
| 0.062 | 0 | 0.004 | 0.022 | 0.020033 | 0.258 | 0.382 | 0.283 | 0.307667 | 0.037861 |
| K63 | | | | | K63 | | | | |
| rep1 | rep2 | rep3 | <χ> | SD | rep1 | rep2 | rep3 | <χ> | SD |
| 0.419 | 0.242 | 0.429 | 0.363333 | 0.060735 | 0.062 | 0 | 0 | 0.020667 | 0.020667 |

**Table S2.** Molar fractions of single tri-Ub isomers in the quaternary mixture treated with various DUBs. Molar fraction values were obtained by deconvolution of the CIU fingerprint of the quaternary mixture with CIU fingerprints of four (K6, K48-, K63-, 6/48-branched) isomers. The mean molar fraction values (<x>) were calculated from three repeats within the same biological sample. Standard deviation was calculated as shown in page S3.

| *+ AMSH* | | | | | *+ OTUB1* | | | | |
|---|---|---|---|---|---|---|---|---|---|
| 6/48-branched | | | | | 6/48-branched | | | | |
| rep1 | rep2 | rep3 | <χ> | SD | rep1 | rep2 | rep3 | <χ> | SD |
| 0.376 | 0.349 | 0.362 | 0.361667 | 0.008373 | 0.067 | 0.039 | 0.073 | 0.059667 | 0.010477 |
| K6 | | | | | K6 | | | | |
| rep1 | rep2 | rep3 | <χ> | SD | rep1 | rep2 | rep3 | <χ> | SD |
| 0.357 | 0.306 | 0.337 | 0.333333 | 0.014836 | 0.546 | 0.551 | 0.532 | 0.543 | 0.005686 |
| K48 | | | | | K48 | | | | |
| rep1 | rep2 | rep3 | <χ> | SD | rep1 | rep2 | rep3 | <χ> | SD |
| 0.267 | 0.292 | 0.286 | 0.281667 | 0.007535 | 0.092 | 0.081 | 0.088 | 0.087 | 0.003215 |
| K63 | | | | | K63 | | | | |
| rep1 | rep2 | rep3 | <χ> | SD | rep1 | rep2 | rep3 | <χ> | SD |
| 0 | 0.053 | 0.0137 | 0.022233 | 0.015884 | 0.2941 | 0.327 | 0.305 | 0.3087 | 0.009676 |
| *+ UCH37* | | | | | *ctrl w/o DUB* | | | | |
| 6/48-branched | | | | | 6/48-branched | | | | |
| rep1 | rep2 | rep3 | <χ> | SD | rep1 | rep2 | rep3 | <χ> | SD |
| 0 | 0.052 | 0.032 | 0.028 | 0.013158 | 0.421 | 0.456 | 0.378 | 0.418333 | 0.023642 |
| K6 | | | | | K6 | | | | |
| rep1 | rep2 | rep3 | <χ> | SD | rep1 | rep2 | rep3 | <χ> | SD |
| 0.389 | 0.416 | 0.419 | 0.408 | 0.009131 | 0.197 | 0.18 | 0.21 | 0.195667 | 0.007545 |
| K48 | | | | | K48 | | | | |
| rep1 | rep2 | rep3 | <χ> | SD | rep1 | rep2 | rep3 | <χ> | SD |
| 0.277 | 0.275 | 0.27 | 0.274 | 0.008675 | 0.171 | 0.15 | 0.167 | 0.162667 | 0.007078 |
| K63 | | | | | K63 | | | | |
| rep1 | rep2 | rep3 | <χ> | SD | rep1 | rep2 | rep3 | <χ> | SD |
| 0.332 | 0.28 | 0.3 | 0.304 | 0.017321 | 0.21 | 0.173 | 0.244 | 0.209 | 0.017769 |

**Table S3.** Molar fractions of single tri-Ub isomers in the quaternary mixture treated with UCH37. The reaction was quenched at the indicated time points, and the molar fractions were calculated by deconvolution of the CIU fingerprint of the quaternary mixtures with CIU fingerprints of four (K6, K48-, K63-, 6/48-branched) isomers at each time point. The mean molar fraction values (<x>) were calculated from three repeats within the same biological sample. Standard deviation was calculated as shown in page S3.

| | 6/48-branched | | | | |
|---|---|---|---|---|---|
| time, min | rep1 | rep2 | rep3 | <χ> | SD |
| 0 | 0.221817 | 0.19354 | 0.15506 | 0.190139 | 0.019346 |
| 15 | 0.109667 | 0.08616 | 0.151331 | 0.115719 | 0.019055 |
| 30 | 0.114074 | 0.118256 | 0.134939 | 0.122423 | 0.006373 |
| 60 | 0.107702 | 0.077449 | 0.095904 | 0.093685 | 0.008804 |
| 240 | 0.001158 | 0.028912 | 0.024147 | 0.017313 | 0.009944 |
| | K6 | | | | |
| time, min | rep1 | rep2 | rep3 | <χ> | SD |
| 0 | 0.213617 | 0.332441 | 0.277263 | 0.27444 | 0.03433 |
| 15 | 0.342214 | 0.291034 | 0.297831 | 0.31036 | 0.016048 |
| 30 | 0.289789 | 0.320435 | 0.244852 | 0.285025 | 0.021948 |
| 60 | 0.380341 | 0.268943 | 0.279604 | 0.309629 | 0.035489 |
| 240 | 0.35535 | 0.415044 | 0.346741 | 0.375583 | 0.022173 |
| | K48 | | | | |
| time, min | rep1 | rep2 | rep3 | <χ> | SD |
| 0 | 0.296716 | 0.325228 | 0.336451 | 0.319465 | 0.011827 |
| 15 | 0.273682 | 0.254691 | 0.207486 | 0.245286 | 0.019679 |
| 30 | 0.214948 | 0.213623 | 0.165439 | 0.198003 | 0.016287 |
| 60 | 0.303346 | 0.18348 | 0.273469 | 0.253431 | 0.036023 |
| 240 | 0.334792 | 0.376853 | 0.354329 | 0.355449 | 0.014873 |
| | K63 | | | | |
| time, min | rep1 | rep2 | rep3 | <χ> | SD |
| 0 | 0.26785 | 0.148791 | 0.231226 | 0.215956 | 0.035207 |
| 15 | 0.274437 | 0.368115 | 0.343352 | 0.328635 | 0.028026 |
| 30 | 0.381189 | 0.336142 | 0.454771 | 0.390701 | 0.034574 |
| 60 | 0.208611 | 0.470128 | 0.351022 | 0.343254 | 0.075593 |
| 240 | 0.3087 | 0.179191 | 0.274783 | 0.251655 | 0.046112 |

**Table S4.** Molar fractions of branched tri-Ub isomers in the ternary mixture treated with UCH37·Rpn13. The reaction was quenched at the indicated time points, and the molar fraction values were obtained by deconvolution of the CIU fingerprint of the ternary mixture with CIU fingerprints of all branched trimers (K6/K48, K11/K48-, K48/K63-) at each time point. The mean molar fraction values (<x>) were calculated from three repeats within the same biological sample. The normalized mean value was obtained by normalization of the mean value on the total amount of trimer fraction left in the mixture. Standard deviation was calculated as shown in page S3.

| | 6/48-branched | | | 11/48-branched | | | 48/63-brabched | | |
|---|---|---|---|---|---|---|---|---|---|
| time, min | <χ> | <χ>, norm. | SD | <χ> | <χ>, norm. | SD | <χ> | <χ>, norm. | SD |
| 0 | 0.2905 | 13.53527 | 0.142661 | 0.3585 | 16.70359 | 0.523264 | 0.35 | 16.30755 | 0.586128 |
| 1 | 0.32525 | 15.10038 | 0.081661 | 0.222 | 10.30679 | 0.235591 | 0.45125 | 20.95018 | 0.263015 |
| 3 | 0.22575 | 10.01156 | 0.593077 | 0.36775 | 16.30898 | 0.945813 | 0.405 | 17.96094 | 0.37761 |
| 5 | 0.2225 | 8.04471 | 0.703998 | 0.4005 | 14.48048 | 0.326657 | 0.37425 | 13.53138 | 0.97995 |
| 15 | 0.1305 | 3.724992 | 0.849334 | 0.44725 | 12.7663 | 1.135532 | 0.4205 | 12.00275 | 0.428279 |
| 30 | 0 | 0 | 0 | 0.65 | 16.01145 | 0.341325 | 0.349 | 8.596917 | 0.341325 |
| 60 | 0 | 0 | 0 | 0.6855 | 11.85846 | 0.25505 | 0.311 | 5.379989 | 0.276649 |
| 120 | 0.00375 | 0.040406 | 0.049487 | 0.77225 | 8.320994 | 0.068227 | 0.22125 | 2.383969 | 0.032937 |

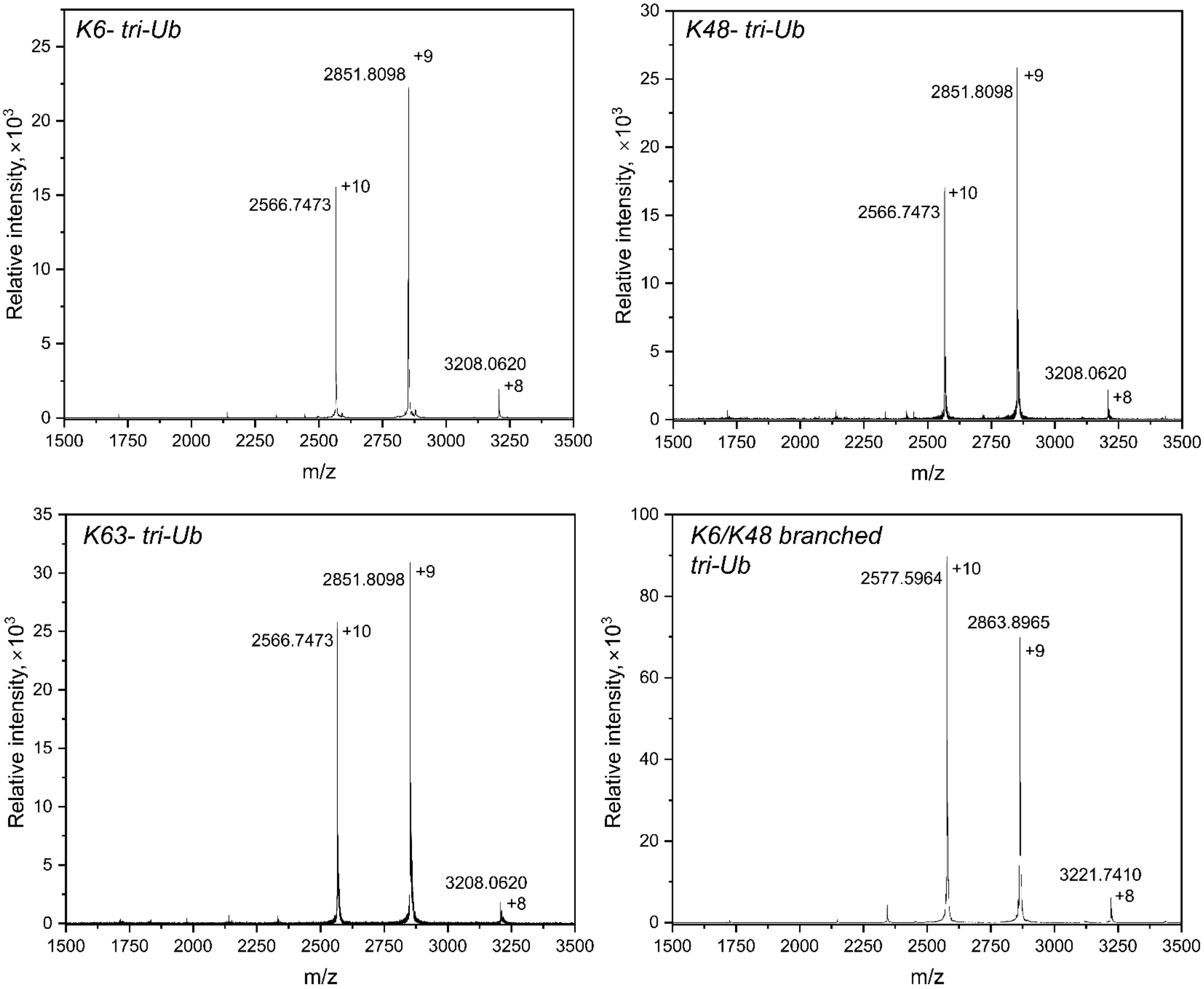


**Figure S1.** MS1 spectra of tri-Ub isomers under native conditions (150 mM ammonium acetate, pH 6.8).

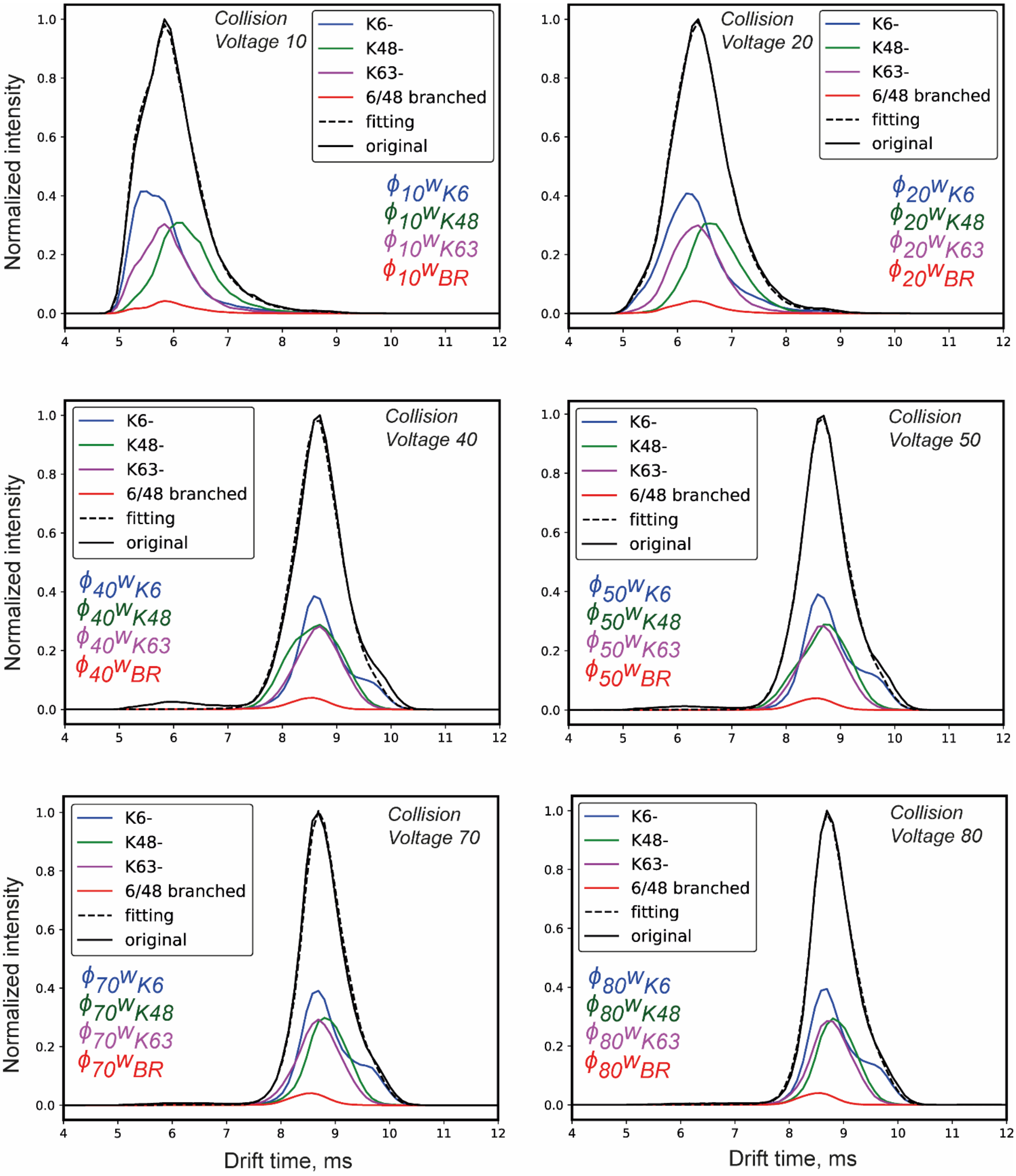


**Figure S2.** Robustness of deconvolution approach using a ternary mixture containing K6-, K48-, and K63-trimers (the K6/K48 branched trimer is missing). The whole CIU fingerprint of the ternary mixture was deconvoluted using the CIU fingerprints of all four single isomers including the branched trimer. Deconvolution was performed at different collision energies. The blue, green, magenta, and red spectra represent the IM-MS spectra of single isomers. The solid black line represents the experimental IM-MS spectrum of ternary mixture. The black dash line corresponds to the IM-MS spectrum obtained from the deconvolution algorithm.

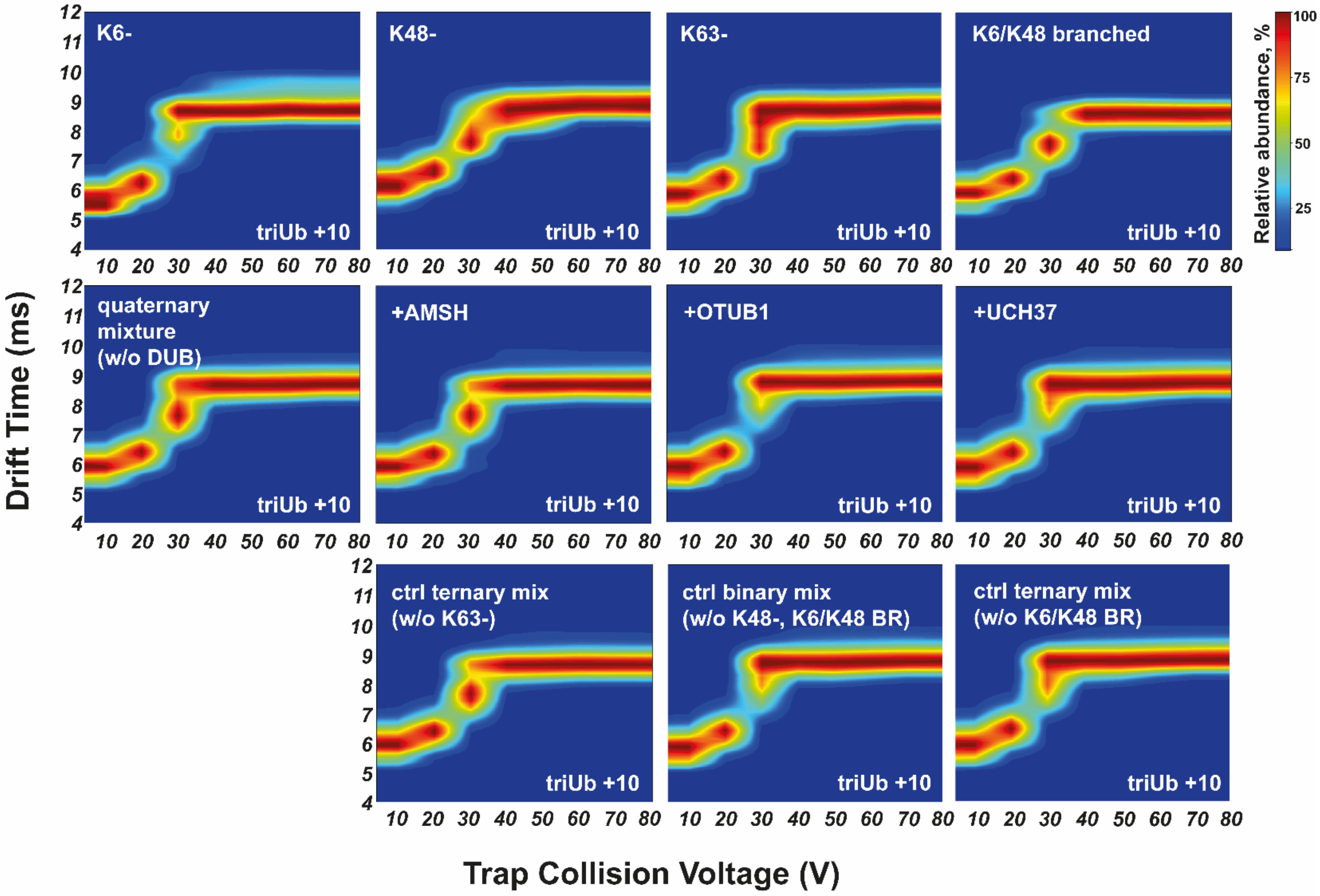


**Figure S3.** CIU fingerprints of quaternary mixtures in the presence and absence of different DUBs. All CIU fingerprints were averaged between three repeats. *Top panel:* CIU fingerprints of individual isomers (from left to right: K6-, K48-, K63-, K6/K48 branched). *Middle panel:* CIU fingerprint of the quaternary mixture before and after cleavage with different DUBs (from left to right: before DUB, +AMSH, +OTUB1, +UCH37). *Bottom panel:* CIU fingerprints of the control ternary mixtures missing specific trimers.

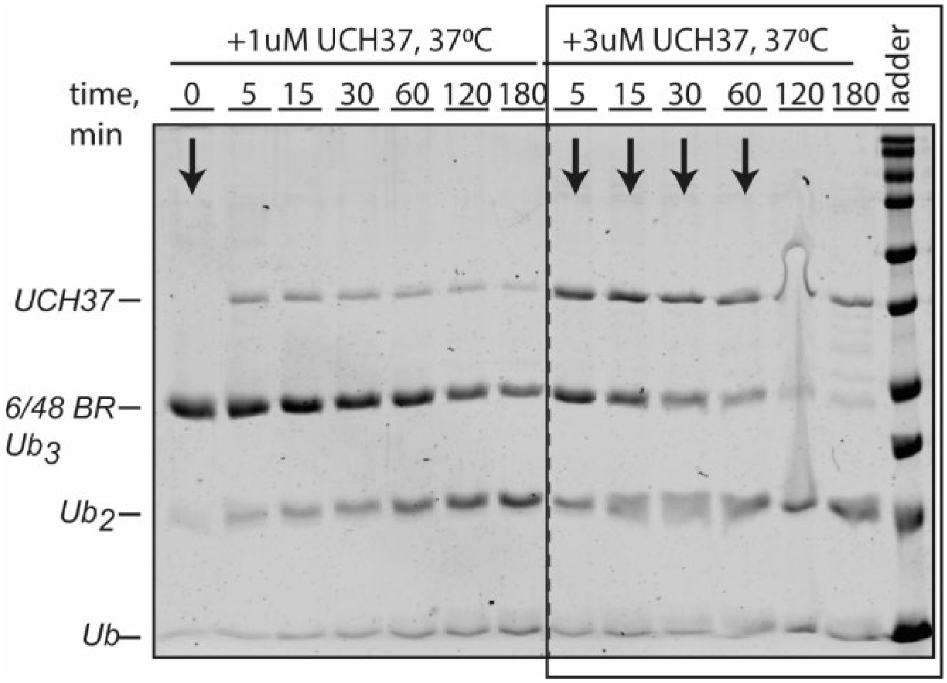


**Figure S4.** UCH37/UCHL5-dependent cleavage of the K6/K48-branched trimer. 10uM of 6/48-branched trimer was treated with 1 uM or 3 uM of UCH37/UCHL5, and the reaction was quenched at the indicated time points. The black rectangle highlights the reaction conditions that were chosen for the IM-MS-based time-dependent DUB assay described in the main text (page 12, Figure 5). Black arrows represent the time points (240 min time point is not shown on the gel) chosen for the IM-MS-based analysis.

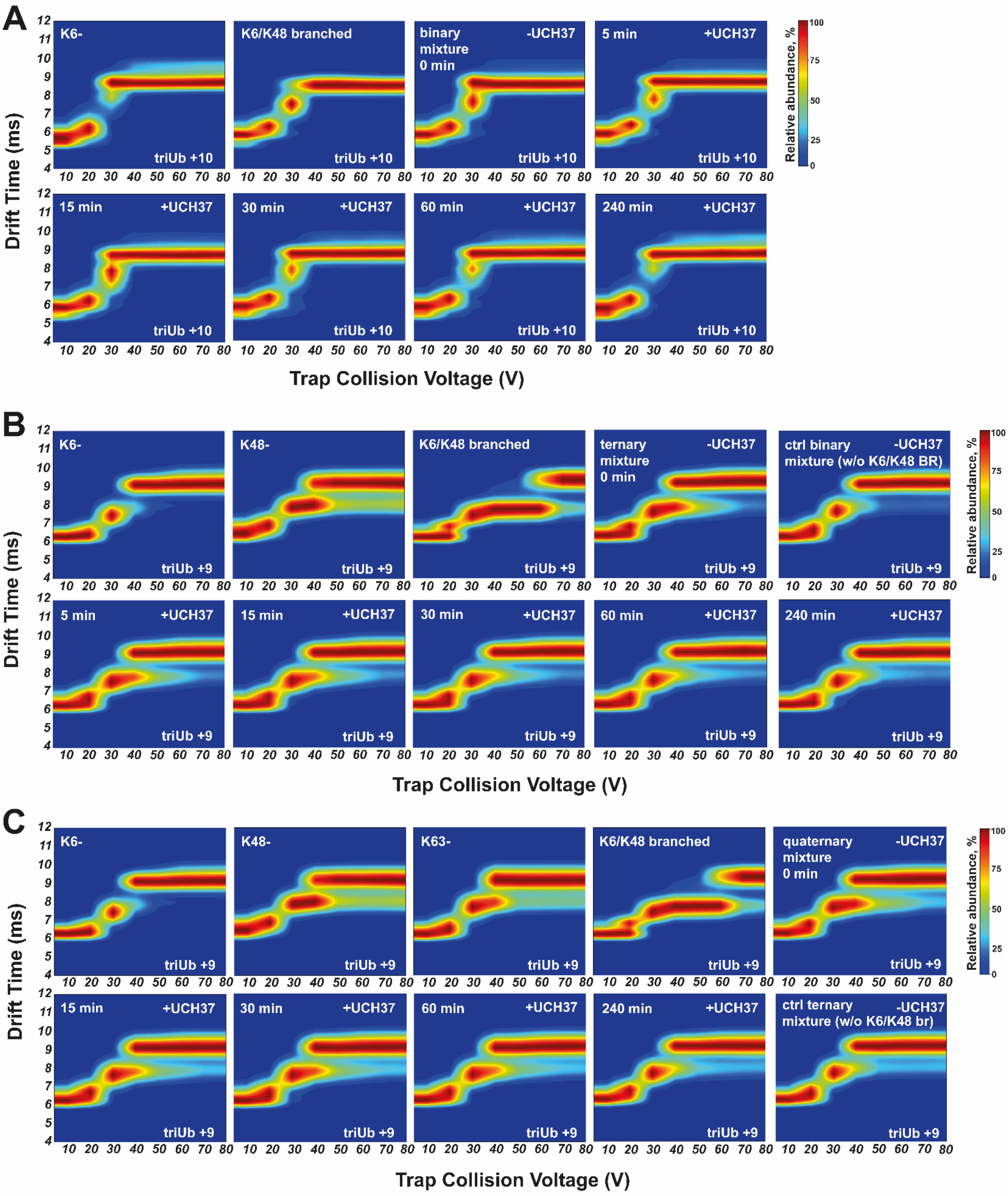


**Figure S5.** CIU fingerprints were obtained from the time-course degradation assays of binary, ternary, or quaternary mixtures with UCH37/UCHL5. All CIU fingerprints were averaged between three repeats. A) Time-dependent degradation of the binary mixture, consisting of K6- and K6/K48 branched trimers; B) Time-dependent degradation of the ternary mixture, consisting of K6-, K48-, and K6/K48 branched trimers; C) Time-dependent degradation of the quaternary mixture, consisting of K6-, K48-, K63-, and K6/K48 branched trimers. Control mixtures represent the mixtures that are expected to form after UCH37/UCHL5 cleavage and were prepared by mixing the specific isomers in an equimolar ratio to compare with the data obtained after actual cleavage.

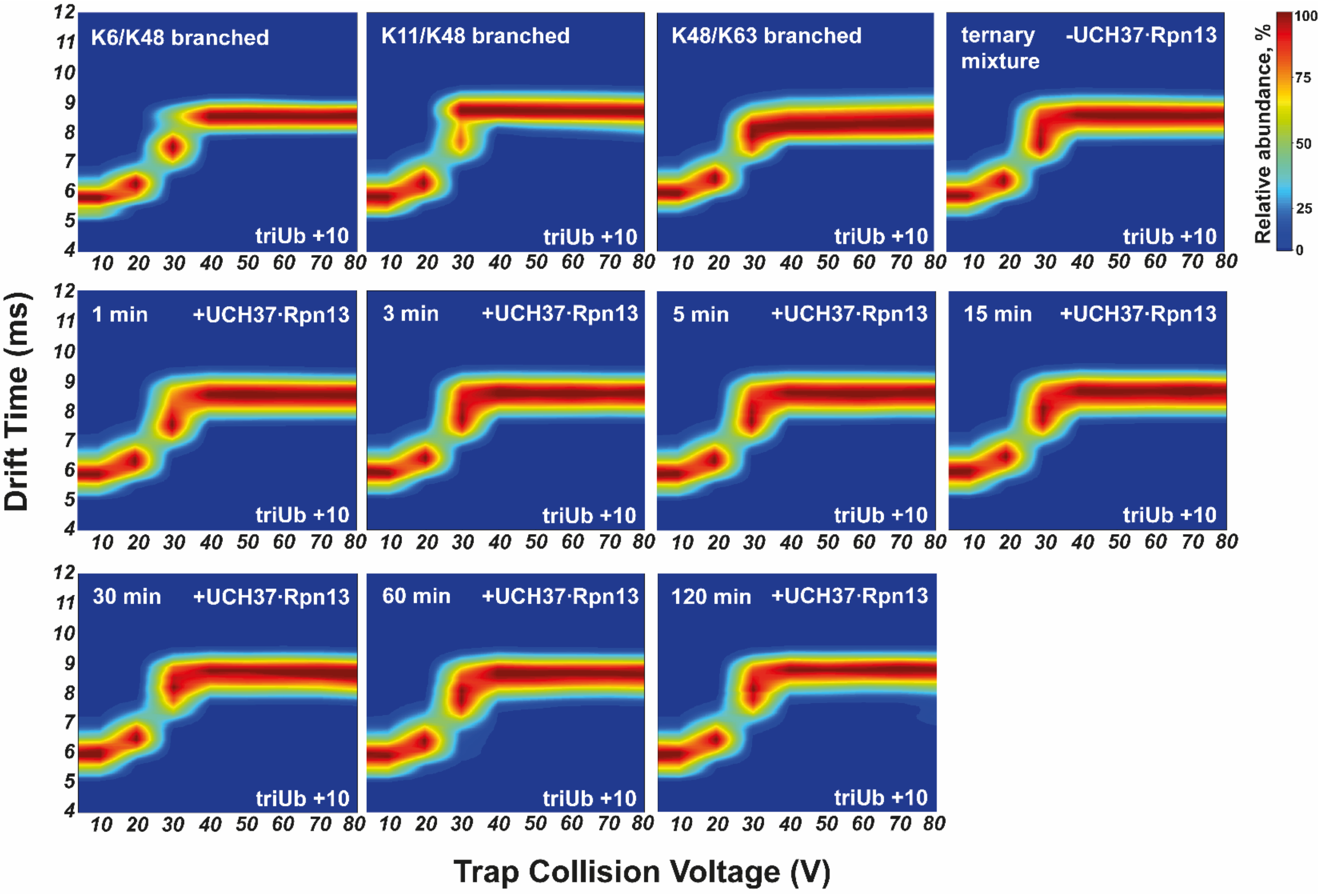


**Figure S6.** CIU fingerprints were obtained from the time-course degradation assays of the ternary mixture, consisting of K6/K48, K11/K48, and K48/K63- branched trimers with UCH37·Rpn13 complex. All CIU fingerprints were averaged between three repeats.

**Protein expression/purification**

*E1, UBE2R1, and UBE2N/UBE2V2* were expressed in Rosetta 2(DE3)pLysS E.coli cells in LB media supplemented with appropriate antibiotics at 37°C to $OD_{600}$ 0.6-0.8 and after induction with IPTG incubated at 18°C for 16 h. Cultures were harvested, resuspended in lysis buffer A (50 mM Tris pH 7.5, 300 mM NaCl, 1 mM TCEP and 10 mM imidazole), lysed by sonication, and clarified by centrifugation. Clarified lysate was then incubated with Ni-NTA resin for 2 h, washed with lysis buffer A, and eluted into Ni-NTA elution buffer (lysis buffer A plus 300 mM imidazole).

*Ube2L3 and UBE2S-UBD* constructs were expressed constructs were expressed in Rosetta2(DE3)pLysS E.coli cells in LB media supplemented with appropriate antibiotics at 37°C to $OD_{600}$ 0.6-0.8 and transferred to 16°C for 16 h after induction with IPTG. Cultures were harvested, resuspended in lysis buffer B (270 mM sucrose, 50 mM Tris pH 8.0, 50 mM NaF, and 1 mM DTT), lysed by sonication, and clarified by centrifugation. Clarified lysate was then incubated with GST resin for 2 h, washed with high salt buffer (25 mM Tris pH 8.0, 500 mM NaCl, and 5 mM DTT) and then by low salt buffer (25 mM Tris pH 8.0, 150 mM NaCl, and 5 mM DTT), and resuspended in 3C protease buffer (50 mM Tris pH 8.0 and 150 mM NaCl) for on-resin cleavage with HRV 3C protease overnight or thrombin cleavage buffer (30 mM Tris pH 7.5, 150 mM NaCl, 2.5 mM CaCl2) for on resin thrombin cleavage.

*NleL (aa 170-782)* was expressed in BL21(DE3)pLysS E.coli cells in LB media supplemented with appropriate antibiotics at 37°C to $OD_{600}$ 0.6-0.8 and transferred to 16°C for 16 h after induction with IPTG. Cultures were harvested, resuspended in lysis buffer C (50 mM Tris pH 8.0, 200 mM NaCl, 1 mM EDTA and 1 mM DTT), lysed by sonication, and clarified by centrifugation. Clarified lysate was then incubated with GST resin for 2 h, washed with lysis buffer, and eluted into GST elution buffer (lysis buffer C plus 10 mM reduced glutathione). Eluate was concentrated in TEV protease buffer (50 mM Tris pH 8.0, 150 mM NaCl, and 0.5 mM TCEP), cleaved overnight with TEV protease.

All enzymes were further purified using size exclusion chromatography on Superdex 200 (GE) or Superdex 75 (GE) columns.

**Deubiquitinating enzymes expression and purification.**

*OTUB1* was expressed in Rosetta 2(DE3)pLysS E.coli cells in LB media supplemented with appropriate antibiotics at 37°C to $OD_{600}$ 0.6-0.8 and transferred to 18°C for 16 h after induction with IPTG. Cultures were harvested, resuspended in lysis buffer A, lysed by sonication, and clarified by centrifugation. Clarified lysate was then incubated with Ni-NTA resin for 2 h, washed with lysis buffer, and eluted into Ni-NTA elution buffer.

*AMSH* was expressed in BL21(DE3)pLysS E.coli cells in LB media supplemented with appropriate antibiotics at 37°C to $OD_{600}$ 0.6 and transferred to 16°C for 16 h after induction with IPTG and addition of ZnCl2. Cultures were harvested, resuspended in lysis buffer D (50 mM Tris pH 7.5, 300 mM NaCl and 2 mM DTT), lysed by sonication, and clarified by centrifugation. Clarified lysate was then incubated with GST resin for 2 h, washed with lysis buffer D, followed by low salt buffer D, and resuspended in 3C protease buffer for on-resin cleavage with HRV 3C protease overnight.

**Purification of UCH37·Rpn13 complex.** UCH37 and RPN13 constructs were expressed in BL21(DE3)pLysS E. coli cells grown at 37°C in LB media supplemented with appropriate antibiotic. Once cells reached an $OD_{600}$ of 0.6, IPTG was added, and the temperature was reduced to 20°C. Cultures were harvested after 16 hr and frozen at −80°C. Cell pellets from separate UCH37 and RPN13 expressions were mixed 1:1 and resuspended in amylose lysis buffer E (50 mM HEPES pH 7.4, 150 mM NaCl, 1 mM EDTA, and 1 mM TCEP), lysed by sonication, and clarified by centrifugation. UCH37•RPN13 were purified following different chromatographic steps. For UCH37•RPN13, clarified lysate was incubated with amylose resin for 2 hr at 4 °C, washed with amylose lysis buffer, and eluted into amylose elution buffer (lysis buffer E plus 10 mM maltose), and incubated overnight with TEV protease at 4°C. Then TEV protease cleaved product was incubated with Ni-NTA resin for 1 hr, washed with Ni-NTA lysis buffer, and eluted with Ni-NTA elution buffer (Ni-NTA lysis buffer plus 300 mM imidazole). The eluate was concentrated and loaded onto a Superdex 200 (GE) gel filtration column in gel filtration buffer (50 mM HEPES pH 7.5, 50 mM NaCl, 1 mM EDTA, and 1 mM TCEP). For UCH37 clarified lysate was subjected to Ni-NTA chromatography followed by further purification using anion exchange chromatography and Superdex 75 (GE) gel filtration column in gel filtration buffer.
